# RE-1 silencing transcription factor is reduced in endometriosis and uterine deletion in mice alters progesterone responsiveness

**DOI:** 10.64898/2026.05.30.728827

**Authors:** Paige Minchella, Ayushi Vashisht, Riley Peterson, Amanda Graham, Sumedha Gunewardena, Wei Cui, Austin Findley, Julie A. Christianson, Vargheese Chennathukuzhi, Warren B. Nothnick

## Abstract

Endometriosis is a steroid-dependent gynecologic disease characterized by progesterone (P4) resistance, subfertility/infertility, and pelvic pain; however, the molecular mechanisms underlying impaired P4 responsiveness in endometriosis tissue are not fully understood. RE-1 silencing transcription factor (REST), a transcriptional regulator implicated in steroid hormone signaling, has emerged as a potential mediator of P4 responsiveness. Here, we investigated the role of REST in endometriosis using human tissues and a uterine-specific Rest conditional knockout mouse model. Immunohistochemical analysis of eutopic endometrium and ectopic lesions from patients with endometriosis revealed significantly reduced nuclear REST expression compared with control endometrium, suggesting loss of functional REST in disease. To assess the physiological consequences of REST deficiency, uterine-specific Rest knockout (Rest ^d/d^) mice were generated. Rest ^d/d^ females exhibited progressive subfertility and hyper-estrogenic uterine tissue characteristics that displayed a blunted responsiveness to P4 treatment. Loss of Rest selectively altered expression of P4-responsive genes associated with endometriosis pathology, despite preserved P4 receptor expression. Following induction of experimental endometriosis, female mice that developed endometriotic-like lesions using Rest-deficient donor tissue developed significantly larger lesions that were less responsive to P4 treatment compared to lesions induced using control tissue. Mechanical sensitivity was modestly increased in mice receiving Rest-deficient tissue, whereas vaginal hyperalgesia was unaffected. These findings identify loss of nuclear REST as a feature of endometriosis and support a role of REST in subfertility, lesion progression, and blunted response to P4. REST may represent a novel molecular contributor to altered P4 responsiveness and a potential therapeutic target in endometriosis.

**Significance Statement:** Endometriosis is a common disease in women characterized by altered steroid hormone signaling, infertility, and pelvic pain. RE-1 silencing transcription factor (REST) is a candidate regulator of steroid hormone signaling in gynecologic disease but a role in endometriosis pathophysiology remains unexplored. To fill this knowledge gap, our study utilizes human endometrial and endometriotic tissues coupled with a conditional knockout mouse model for uterine Rest deficiency. We show that REST is significantly reduced in eutopic and ectopic endometrial tissue from women with endometriosis and that deletion from mouse uterine tissue recapitulates clinical characteristics in women with endometriosis including progesterone resistance, sub-fertility and pelvic pain. These findings will further guide future research to understand impaired steroid signaling in the pathophysiology of endometriosis.

## Introduction

### Endometriosis and progesterone resistance

Endometriosis is a complex, estrogen-dependent gynecological disease that affects approximately 10% of reproductive-age women, or 190 million worldwide, not including the estimated 6 million who are suspected to go undiagnosed.^1,2^ The disease is characterized by the ectopic presence of uterine-like tissue containing endometrial stroma and glands outside of the uterus which can develop into superficial or deep infiltrating lesions and disrupt normal tissue physiology. Aberrant steroid hormone signaling, specifically progesterone (P4) resistance, is implicated in the advancement of disease pathophysiology.

Patients with endometriosis exhibit altered expression levels of several endometrial P4 target genes^3–8^ which may be due to abnormal expression and/or function of P4 receptors (PGR), PGR-A and PGR-B,^9–11^ as well as inflammation, genetics, and epigenetics.^12–15^ The inability of eutopic endometrium and ectopic endometriotic tissue to properly respond to P4, leading to hyper-estrogen action and inflammation, promotes proliferation and survival of the ectopic tissue and the onset of endometriosis-associated symptoms.

Clinical manifestations of the disease that cause many patients to seek medical evaluation include infertility and pelvic pain.^16–19^ For endometriosis-associated infertility, poor oocyte quality, impaired embryo implantation, and P4 resistance are implicated as contributing factors, yet the precise mechanisms remain poorly understood.^12,13,20,21^ In patients with endometriosis, pelvic pain does not always correlate with stage or severity of the disease.^22,23^ However, aberrant P4 response and concurrent overexpression of estrogen receptor-β (ESR2) results in excessive inflammation and sensitization.^8,24,25^ This pro-inflammatory environment brought on by P4 resistance in the tissue could contribute to manifestation of pain symptoms experienced by patients.

P4 resistance is a central mechanism in endometriosis and can influence endometriosis-associated fertility challenges and pelvic pain through promotion of aberrant steroid hormone signaling and increased estrogen activity. These characteristics can also drive lesion survival through blunted P4 responsiveness and inflammation. However, the exact molecular mechanisms and pathways leading to P4 resistance in disease tissue have yet to be fully elucidated. As such, a critical gap in knowledge remains as to why eutopic endometrium and ectopic endometrial-like tissue from patients with endometriosis exhibit altered P4 responsiveness.

### RE-1 silencing transcription factor (REST) and P4 resistance

Although P4 resistance is well documented in endometriosis pathophysiology, the molecular mechanisms driving physiological changes in the tissue that promote an altered P4 responses are not fully understood, limiting the identification of potential new biological targets for treatment. Recent evidence suggests the neuron-restrictive silencer factor/RE1-silencing transcription factor (NRSF/REST) may play a crucial role in steroid hormone responsiveness in reproductive tissue and the transcriptional regulation of P4 target genes, contributing to uterine pathology.^26–28^

REST is a transcriptional repressor that inhibits neuronal gene expression in non-neuronal tissues but can also act as an enhancer of gene transcription.^29,30^ REST functions in the nucleus and binds to highly conserved RE1 elements in target genes.^31,32^ Recent ChIP-seq data support a direct relationship between REST and PGR signaling with observed RE1 sites in roughly 200 REST target genes in close proximity to PGR-A and PGR-B binding sites in human endometrial stromal cells.^27,33^ The loss of functional REST protein in uterine leiomyoma has been associated with an abnormal response to steroid hormones, enhanced estrogen signaling, and a disruption in PGR signaling resulting in altered expression of genes containing both REST and PGR target sequences. Furthermore, coimmunoprecipitation experiments revealed interaction between REST and PGR, and genes containing both REST and PGR target sequences showed aberrant expression in a REST knockout mouse model,^27^ suggesting an epigenetic influence of REST in the regulation of P4 target genes that share both REST and P4 response elements. Additionally, altered expression of these REST-PGR target genes coincided with enhanced ER-α (ESR1) signaling, which supports an overall dysfunctional sex steroid regulation in the tissue.^26^

The Chennathukuzhi lab^27,34^ previously reported that the loss of REST protein, but not mRNA, in uterine leiomyoma promotes aberrant expression of REST and P4 target genes.^21,27,34^ Beyond the scope of uterine pathology, REST also acts as a tumor suppressor in mammary epithelial cells and is shown to be down-regulated in breast cancer, lung cancer, and colon cancer.^5–7,9^ These findings support a model of REST as a key modulator in steroid hormone signaling, with presence of REST necessary for full P4 responsiveness. Whether a similar REST-dependent mechanism contributes to P4 resistance in endometriosis remains unexplored. Our previously published RNA-sequencing data compared differentially expressed uterine mRNAs of control and *Rest* conditional knockout mice during the diestrus stage of the estrus cycle. Interestingly, of the hundreds of differentially expressed transcripts between genotypes, numerous P4 target genes important for endometrial function displayed reduced expression including *Foxo1*, *Irs2* and *Igfbp1* while factors involved in neurogenesis including brain-derived neurotropic factor *(Bdnf)* were elevated in our mouse model. These patterns of expression parallel those in humans where expression of *FOXO1*, *IRS2* and *IGFBP1* is reduced in endometrial tissue from patients with endometriosis and P4 resistance,^7^ while endometrial *BDNF* expression is elevated in endometriosis.^27,34–38^

Collectively, these data support a model in which reduction of REST protein leads to altered steroid hormone responsiveness and support REST as a suitable candidate in P4 resistance in endometriosis. We first aimed to characterize the expression of REST protein in endometriosis utilizing a robust, well-characterized number of human specimens. Next, we investigated how this reduction in uterine REST protein, as seen in uterine leiomyoma and breast cancer, contributes negatively to fertility and pelvic pain symptomology often associated with endometriosis using a novel Rest-deficient mouse model. Additionally, we investigated loss of REST protein in endometriotic-like lesion survival *in vivo*. These studies are an essential first step to determine whether REST/regulation of REST could be examined as a potential therapeutic target for endometriosis treatment.

## Results

### Nuclear REST protein expression is reduced in patients with endometriosis

Previous studies have demonstrated aberrant expression of REST in human cell lines and tissue, implicating REST in steroid hormone signaling and tumor pathogenesis including breast cancer and uterine leiomyoma. Currently, there are no published studies that have investigated the influence of REST expression in the setting of endometriosis, a disease characterized as being steroid-dependent. We first aimed to characterize the subcellular localization and expression pattern of REST protein in human endometriosis by immunohistochemical assessment of ectopic endometriotic lesion tissue and matched eutopic endometrium from patients with confirmed endometriosis (n = 16) and then compared REST expression to eutopic control tissue from patients without endometriosis (n = 15) (**Table 1**).

**Table 1.**
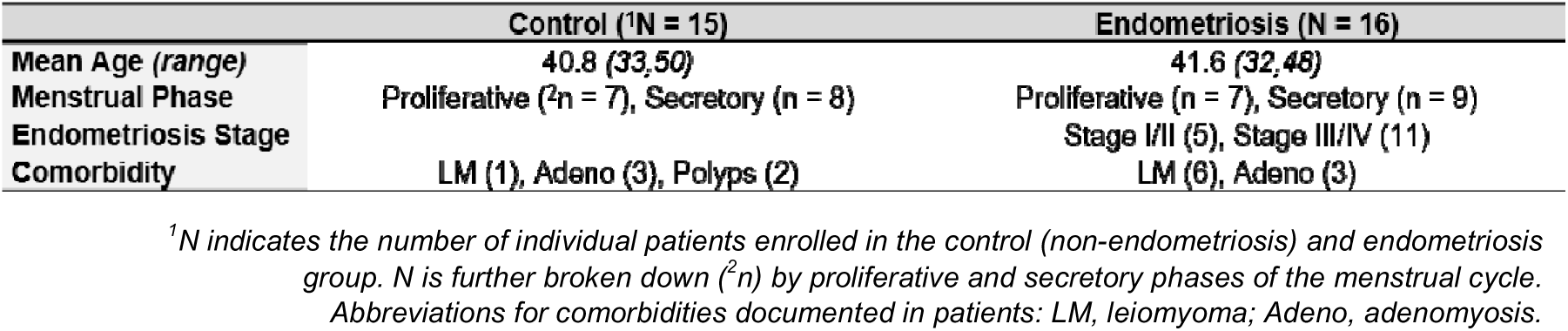
Patient demographics.

Immunohistochemical analysis revealed predominant nuclear localization of REST protein expression in glandular epithelium and endometrial stroma of eutopic endometrium tissue collected from control patients (**black arrows, Figure 1A, F**) during both proliferative and secretory stages of the menstrual cycle. In contrast, nuclear REST expression in glandular epithelium and stroma was reduced or absent in ectopic lesion and eutopic endometrial tissue from patients with endometriosis, independent of menstrual cycle stage (**Figure 1B-C, G-H**). Quantitative HSCORE (Histological Score) analysis confirmed that nuclear REST expression was significantly lower in ectopic endometriosis lesion tissue *(Glands: proliferative, p = 0.0002; secretory, p = 0.0377. Stroma: proliferative, p < 0.0001; secretory, p = 0.0011)* and matched eutopic endometriosis endometrium *(Glands: proliferative, p < 0.0001; secretory, p = 0.0001. Stroma: proliferative, p < 0.0001; secretory, p < 0.0001)* as compared to nuclear REST expression in the glandular epithelium and stroma of control eutopic endometrium (**Figure 1D-E, I-J**). Additionally, in control samples, nuclear REST expression was significantly higher compared to cytoplasmic expression in both the glandular and stromal *(Glands: proliferative, p = 0.0306; secretory, p = 0.0311. Stroma: proliferative, p < 0.0001; secretory, p = 0.0006)*, unlike in endometriosis eutopic endometrium where nuclear REST expression was significantly reduced relative to cytoplasmic in glands alone *(proliferative, p = 0.0158; secretory, p = 0.0134)*. Nuclear REST expression in ectopic endometriotic lesion tissue of glands *(proliferative, p = 0.0656; secretory, p = 0.0568)* and eutopic *(proliferative, p = 0.1495; secretory, p = 0.2459)* and ectopic *(proliferative, p = 0.2020; secretory, p = 0.6329)* tissue in stroma trended lower compared to cytoplasmic expression but overall was not significant. No differences were observed between proliferative and secretory phases of the menstrual cycle within either group. These findings demonstrate abnormal subcellular localization with reduced nuclear expression suggesting loss of functional REST as a nuclear transcription factor in endometrial tissue of patients with endometriosis. Variability in REST expression in ectopic endometriotic lesion tissue may reflect differences in disease stage or lesion establishment/maturity. Additional considerations include the location and lesion type.

**Figure 1.**
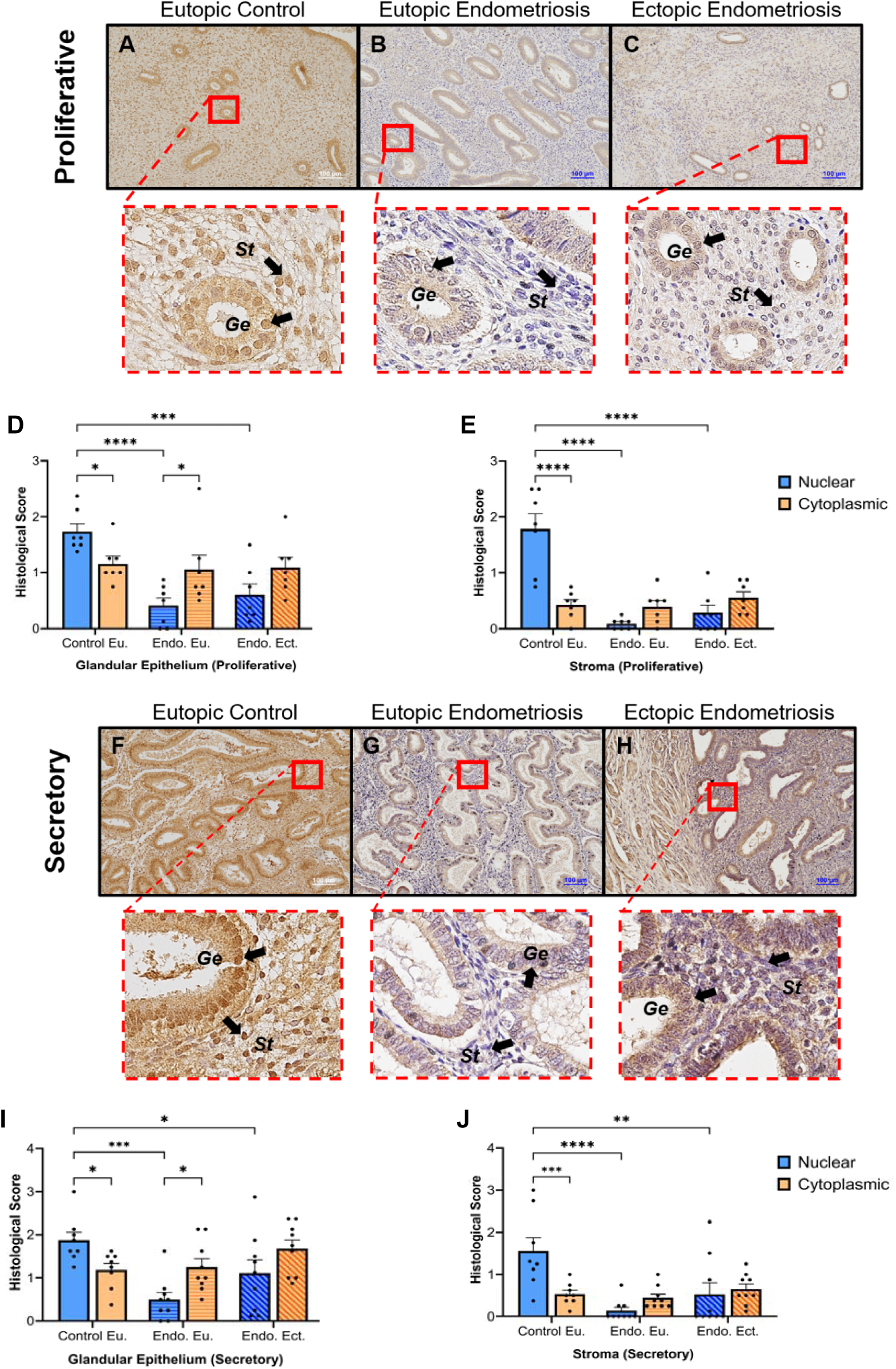
Rest protein expression in human eutopic control, eutopic endometrium from endometriosis, and matched ectopic endometriotic lesion tissue. Representative immunohistochemical images of REST protein expression localization during the proliferative phase (Control, n = 7; Endometriosis, n =7) (**A-C**) and secretory phase (Control, n = 8; Endometriosis, n = 9) (**F-H**) (REST, brown staining; hematoxylin, purple counterstain) from ectopic endometriotic lesion tissue (Ectopic Endometriosis) and matched eutopic endometrial tissue (Eutopic Endometriosis) from patients with confirmed endometriosis, and eutopic control endometrium from patients without endometriosis (Eutopic Control). Images were taken at 10X (scale bar indicates 100μm); dashed-line boxes are the amplified images of the area designated by the solid box region outlined in the respective panel. Control sample (**A, proliferative; F, secretory**), REST was localized to endometrial stroma (**St**) and glandular epithelium (**Ge**), and localization was predominantly nuclear (black arrows). In contrast, nuclear REST protein was low/absent in eutopic (**B proliferative; G, secretory**) and ectopic/lesion tissue (**C, proliferative; H, secretory**) in endometriosis, with REST expression predominantly cytoplasmic. Tissue samples were assessed using H-Score for REST protein expression in proliferative endometrial glandular epithelium (**D**), proliferative endometrial stroma (**E**), secretory endometrial glandular epithelium (**I**), and secretory endometrial stroma (**J**). Samples were scored on a scale of 0-3 based on previously established parameters for histological scores with 0 being little/absent staining and 3 being the most strong/intense staining. H-Scores were analyzed using two-way ANOVA (Tukey’s multiple comparison test) between nuclear and cytoplasm within group, nuclear alone across all groups, and cytoplasm alone across all groups. Glands and stroma were analyzed separately for both secretory and proliferative tissue samples. *Significance is set at *p < 0.05; **p <0.01; ***p < 0.001; ****p < 0.0001*. Abbreviations: Cont. Eu. = Control eutopic endometrium, Endo. Eu. = Endometriosis eutopic endometrium, Endo. Ect. = Endometriotic ectopic lesion tissue. Data is represented as mean ± SEM. Scale bars are 100µm. REST, human and mouse protein; *REST,* human gene symbol; *Rest,* mouse gene symbol.

### Uterine Rest deficient female mice display subfertility

Reduced expression of REST in human endometriosis tissues prompted further investigation of the potential contribution that loss of REST may have to the clinical manifestations of endometriosis. To do this, we created an *in vivo* model of whole uterine REST knockout using the Cre-lox system. *Rest ^fl/fl^* mice were mated to *Pgr ^Cre/+^*mice to generate a *Rest ^fl/fl^ Pgr ^Cre/+^ (Rest ^d/d^*) line, in which the expression of Cre recombinase is under the control of progesterone receptor (PGR) promoter action. The REST gene is flanked by LoxP sites and is conditionally deleted from tissue where and when PGR is expressed. To assess fertility, breeding trials were conducted using mature, 2-month-old female *Rest ^d/d^* mice (n = 8) that were mated with mature wild-type C57BL/6 males of proven fertility. As a control group, *Rest ^fl/fl^*female mice (n = 9) were also mated with mature wild-type C57BL/6 males. After the first round, when pups were weaned from the dams, each mother was re-mated. After 3 rounds of consecutive mating, the total number of litters and pups were calculated for both genotypic groups of females. As shown in **Table 2**, after 3 breeding cycles, *Rest ^fl/fl^* female dams produced a total of 248 pups across 26 litters, with 96% of females having successful parturition and birth of viable offspring. However, *Rest ^d/d^* female dams only produced 57 pups in total across 11 litters, with 46% of females having successful parturition across 3 breeding cycles.

**Table 2.** Number of successful deliveries/litters and pups during breeding trial. The total number of litters and pups (*) overall and (**/***) per round of breeding. Each round of mating was broken down into (**) number of litters per genotype (out of total number of dams per genotype in that round) as some dams did not successfully produce a litter, and (***) total number of pups from all dams in that genotype (average number of pups per dam ± SEM).

|  | **Rest**<sup>fl/fl</sup> | **Rest**<sup>d/d</sup> |
| --- | --- | --- |
| **\*Total Litters** | 26 | 11 |
| **\*Total Pups** | 248 | 57 |
| **\*\*Litters** | Litters (Total Dams) | Litters (Total Dams) |
| Round 1 | 9 (9) | 8 (8) |
| Round 2 | 9 (9) | 1 (8) |
| Round 3 | 8 (9) | 2 (8) |
| **\*\*\*Pups** | Total (Avg±SEM) | Total (Avg±SEM) |
| Round 1 | 72 (8.0±0.6) | 33 (4.12±0.93) |
| Round 2 | 78 (8.67±1.05) | 5 (0.625±0.625) |
| Round 3 | 78 (8.67±1.21) | 13 (1.625±2.8) |

Compared to *Rest ^fl/fl^* control females, *Rest ^d/d^* females exhibited a significant decrease in total number of litters per breeding round and the litter size per dam from the first round of breeding to the subsequent second and third (**Figure 2**). The proportion of dams with successful litters decreased from round 1 to round 2, with only 1 out of 8 dams in round 2 producing pups and remained low in round 3 (2 out of 8 dams, **Table 2**). In contrast, *Rest ^fl/fl^* dams maintained a stable average litter size across all mating rounds (**Table 2**, **Figure 2**). Across all consecutive rounds of mating, *Rest ^d/d^* dams produced significantly smaller litter sizes compared to *Rest ^fl/fl^* dams in the same round (**Figure 2**). Additionally, only 43% of pups from *Rest ^d/d^* females survived until weaning compared to 98% from control. Due to PGR expression in the mammary glands, we investigated if loss of REST affected lactation and feeding in KO females. The average litter weight of the two genotypes (*Rest ^fl/fl^*= 10.22 grams; *Rest ^d/d^* = 9.35 grams) was not found to be statistically significant, however, poor lactation cannot be fully ruled out as a cause of neonatal death in the REST deficient mice. A decline in successful pregnancy and parturition in *Rest ^d/d^* females is consistent with subfertility, suggesting that loss of REST may contribute to impaired fertility. However, whether the loss of REST is associated with development or exacerbation of endometriosis and is a mechanism that contributes to subfertility as a clinical manifestation of the disease remains to be determined.

**Figure 2.**
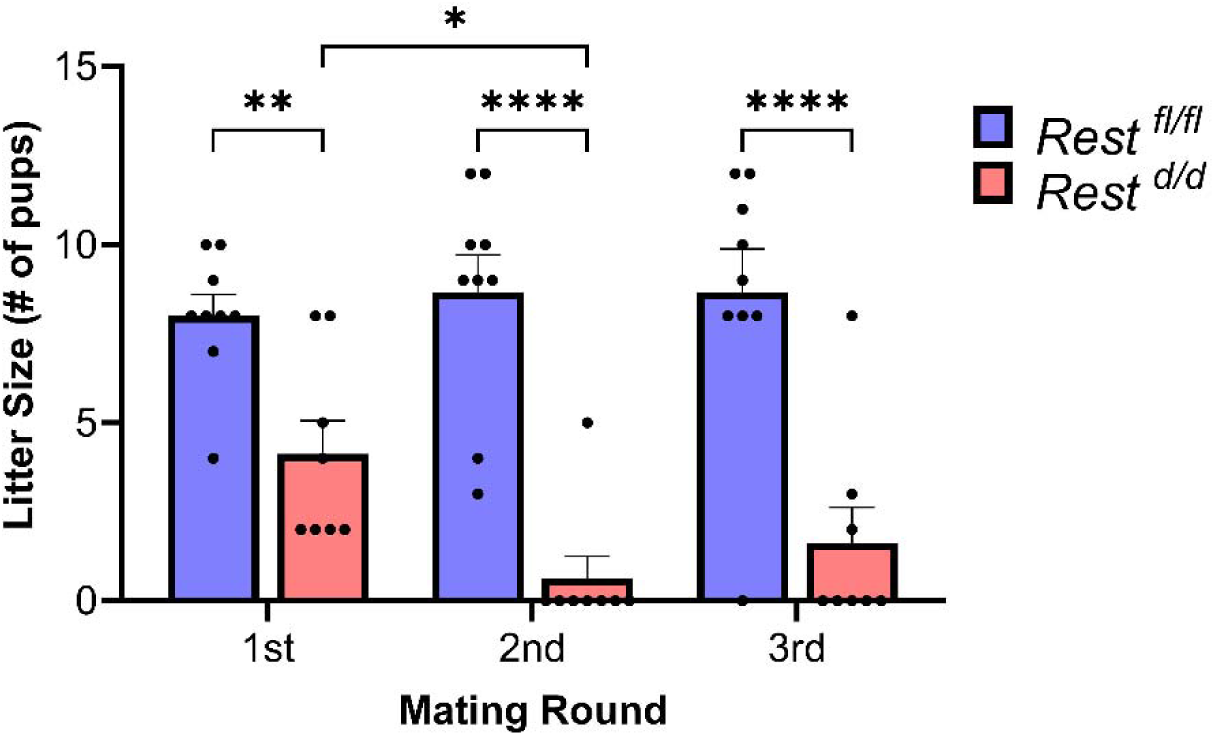
Loss of uterine Rest is associated with subfertility in female mice. 2-3-month-old female mice of control genotype, *Rest ^fl/fl^* (n = 9), and experimental uterine Rest conditional knockout, *Rest ^d/d^* (n = 8), were mated with wild type males of proven fertility. The number of living pups (male and female) in each litter was recorded for each female/genotype (y-axis) after each round of consecutive mating (x-axis), 3 breeding cycles overall. Experimental *Rest ^d/d^* females on average had significantly fewer pups per litter after each round of mating as compared to the control *Rest ^fl/fl^* females. Data for number of pups per litter between genotype within each round of mating and within each genotype across all rounds of mating were analyzed using two-way ANOVA (Tukey’s multiple comparison test). *Significance is set at *p < 0.05; **p <0.01; ***p < 0.001; ****p < 0.0001*. Data is represented as mean ± SEM.

### P4 treatment fails to suppress hyper-estrogen-induced increased uterine wet weight in Rest conditional knockout mice

Gross morphology of *Rest ^d/d^* mouse uteri displayed hallmarks of unopposed estrogen action in the tissue including an increased uterine wet weight and distended, fluid-filled uterine horns as compared to the uteri of control mice (**Figure 3**). Histological assessment using hematoxylin and eosin staining identified enlarged cystic glands in the endometrium of the uterine Rest deficient females. These phenotypic differences in the absence of uterine Rest expression are evidence of a hyper-estrogenic effect that was consistently observed in Rest deficient females aged 6-, 9-, and 12-months-old.

**Figure 3.**
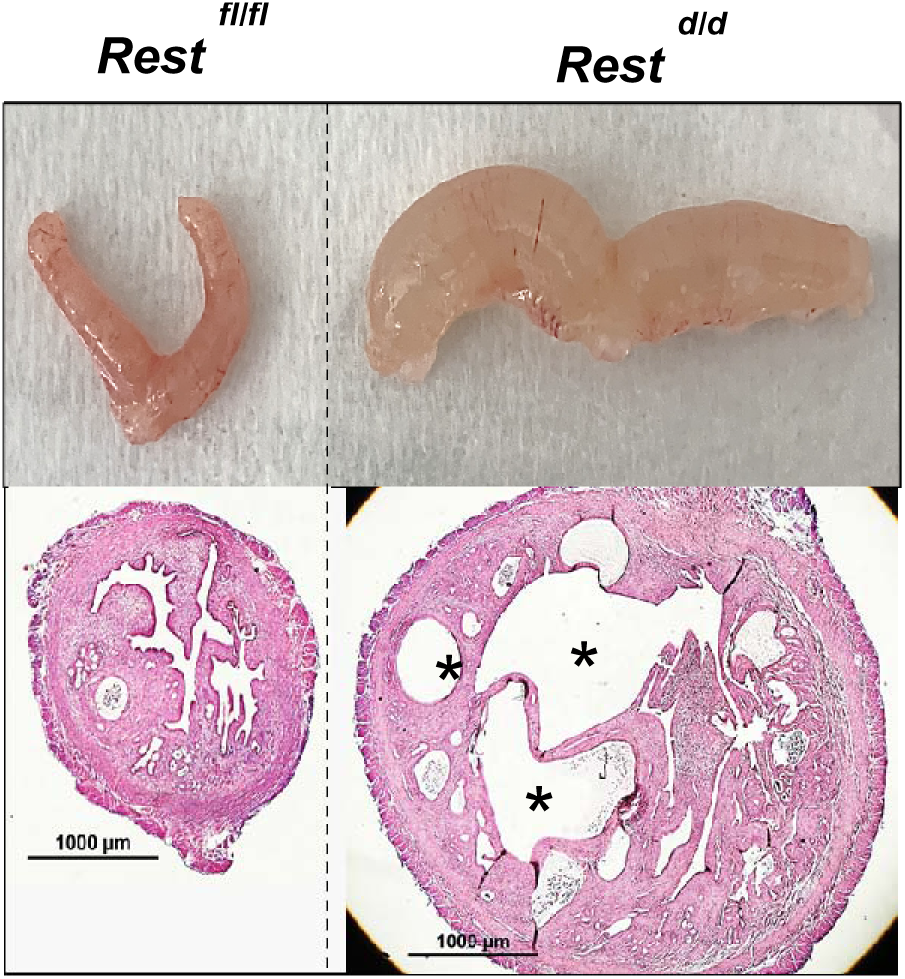
Conditional deletion of uterine Rest expression leads to altered uterine morphology and histology. Uterine tissue was harvested from *Rest ^fl/fl^* and *Rest ^d/d^* mice at 6-, 9-, and 12-months of age and were assessed histologically. *Rest ^d/d^*, but not *Rest ^fl/fl^,* uteri exhibited signs of unopposed estrogen action including increased uterine wet weight, diameter, and large cystic glands (9-month group is shown as a representative image, other age groups not shown; n = 6/genotype/age). Top two panels are *Rest ^fl/fl^* and *Rest ^d/d^* uteri at harvest and the bottom two panels are cross-section H&E immunohistochemical stained images (2X magnification). *cystic gland structures.

Previous studies identified a critical association between REST protein and PGR signaling, indicating that the interaction between the two may contribute to a disruption in steroid hormone response and sensitivity of the tissue.^26,27^ To examine this more closely, we assessed steroid hormone responsiveness in ovariectomized REST deficient *Rest ^d/d^* and control *Rest ^fl/fl^* female mice administered subcutaneous injections of either estradiol-17β (E2) alone or a combination of E2 and P4. Uteri from both genotypes collected at 0hr (no treatment), 6hrs post-injection, and 24hrs post-injection were weighed and expressed as uterine wet weight as a percentage of total body weight per mouse. Similar to results without administration of steroids in **Figure 3**, *Rest ^d/d^* mice exhibited a hyper-estrogenic phenotype and displayed significantly heavier uterine wet weights than *Rest ^fl/fl^* mice following both E2 (**Figure 4A**) treatment at baseline (0hr, no treatment) as well as 6hrs and 24hrs post steroid hormone treatment. Additionally, *Rest ^d/d^* uterine tissue failed to respond to P4, maintaining heavier uterine wet weights as compared to controls following combination treatment of E2 and P4 (**Figure 4B**). To further investigate the blunted P4 response in REST deficient uterine tissue, *Rest ^d/d^* and *Rest ^fl/fl^* female mice were treated chronically with unopposed P4, or vehicle as control, daily for three days before uterine tissue was collected and weighed on the third day (**Figure 4C**). *Rest ^fl/fl^* female mice in both vehicle and treatment groups had a significantly lower uterine wet weight as compared to *Rest ^d/d^*mice, and only *Rest ^fl/fl^* mice displayed a significant decrease in uterine wet weight with P4 treatment compared to vehicle. Chronic, unopposed P4 failed to reduce uterine wet weight in *Rest ^d/d^* mice, with weights remaining elevated in both vehicle and progesterone treated females. Together, these results demonstrate that loss of uterine Rest expression in vivo disrupts P4 responsiveness in the tissue and contributes to a persistent hyper-estrogenic phenotype, suggesting a critical role for Rest in regulating steroid hormone signaling in the uterus.

**Figure 4.**
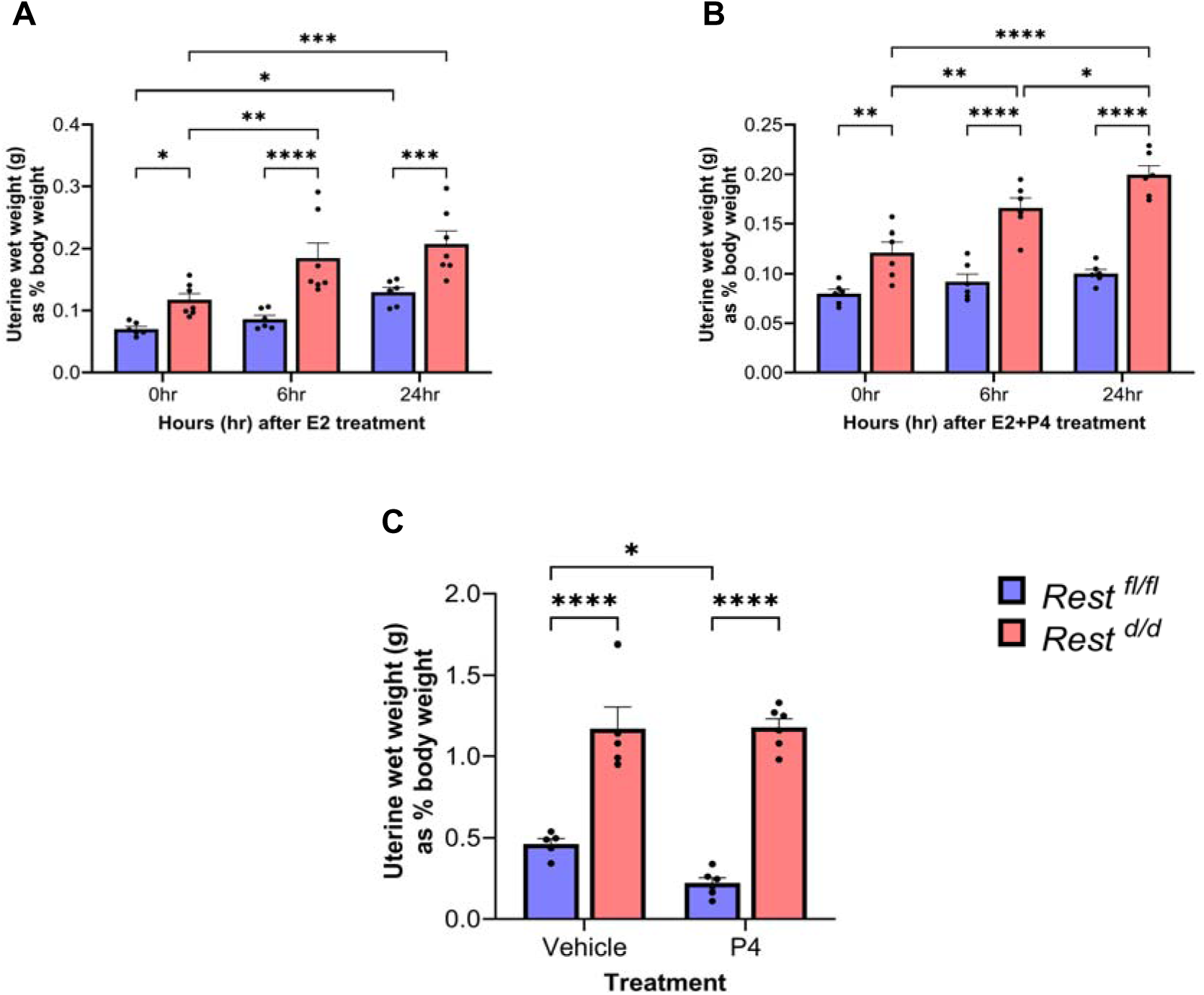
Estrogen treatment reveals hyper-estrogenic effect and failure to suppress estrogen-induced increase in uterine wet weight in Rest conditional knockout mice. Ovariectomized female mice of both genotypes, *Rest ^fl/fl^* and *Rest ^d/d^,* were split into groups and administered subcutaneous injections of (**A**) estradiol-17β (E2) only (*Rest ^fl/fl^*, n = 6/time point; *Rest ^d/d^*, n = 7/time point) or (**B**) combination of progesterone (P4) + E2 (*Rest ^fl/fl^*, n = 6/time point; *Rest ^d/d^*, n = 6/time point), and uteri was harvested at different time points (0hr/no treatment, 6hrs post-injection, and 24hrs post-injection). Uterine wet weight was calculated as a percentage of total body weight per mouse. *Rest ^d/d^* revealed hyper-estrogen effects with significantly heavier uterine wet weights as compared to *Rest ^fl/fl^*at baseline (0hr, no treatment) and following treatment with E2 only in the 6hrs and 24hrs groups. *Rest ^d/d^*, but not *Rest ^fl/fl^,* showed failure to respond to P4 treatments with heavier uterine wet weights at baseline and following P4+E2 treatment. Additionally, in a separate group of mice, chronic treatment of unopposed P4 (**C**) also failed to suppress uterine wet weight in *Rest ^d/d^,* where P4 or vehicle (control) was administered subcutaneously, and only *Rest ^fl/fl^* displayed lower uterine wet weights in the P4 group as compared to vehicle controls, with *Rest ^d/d^* uterine weights significantly higher in both vehicle and P4 treatment groups as compared to *Rest ^fl/fl^* control mice (*Rest ^fl/fl^*: chronic P4, n = 6; vehicle, n = 5. *Rest ^d/d^*: chronic P4, n = 6; vehicle, n = 5). Uterine wet weights as percent body weight were analyzed between genotype within time point and within genotype across time points using two-way ANOVA (Tukey’s multiple comparison test). Data is represented as mean ± SEM. *Significance is set at *p < 0.05; **p <0.01; ***p < 0.001; ****p < 0.0001*.

### Loss of uterine Rest is associated with impaired regulation of select P4 response genes

To determine if this disruption in P4 tissue response extended to P4 signaling and gene transcription downstream, uterine tissue collected from ovariectomized *Rest ^d/d^* and *Rest ^fl/fl^* mice following E2+P4 treatment at 0hr (no treatment), 6hrs post-injection, and 24hrs post-injection was subjected to qRT-PCR to analyze mRNA expression levels of known P4 responsive genes dysregulated in the endometrium of patients with endometriosis.^7^ Progesterone receptor (*Pgr*), a target gene upregulated in the human eutopic endometrium in patients with endometriosis during the early secretory stage, showed a significant increase in expression 6 hours after E2+P4 treatment in both *Rest ^fl/fl^* and *Rest ^d/d^* uterine tissue. Notably, *Rest ^d/d^ Pgr* levels were significantly higher compared to *Rest ^fl/fl^*, with fold-changes of 4.8 and 3.2 respectively (**Figure 5A**), a similar pattern documented in the endometrium of patients with endometriosis as relative to healthy controls. Secreted phosphoprotein-1 (Spp1), also known as osteopontin, a glycoprotein implicated in regulation of the immune response, had significantly lower expression compared to control mice after steroid hormone treatment at 24 hours (8.8- and 19.8-fold-change, respectively), consistent with downregulation in human endometriosis (**Figure 5B**). However, mucin-1 (*Muc1*) expression, while differentially expressed, was elevated in *Rest ^d/d^* compared to controls at 6hrs (8.6- and 2.5-fold-change, respectively) and 24hrs (7.5- and 1.1-fold-change, respectively) after treatment (**Figure 5C**), differing from the reduction seen in human endometriosis. Loss of uterine REST was also associated with an increase in matrix metalloproteinase-9 (*Mmp9*) at 24 hrs (*Rest ^fl/fl^* fold-change of 0.66, *Rest ^d/d^* fold-change of 1.65) and a sustained elevation of matrix metalloproteinase-24 (*Mmp24*) across all time points in *Rest ^d/d^* compared to *Rest ^fl/fl^*(150-to-200-fold-change range) (**Figure 5D-E**). Some P4 responsive genes reported to be differentially expressed in human endometriosis are not impaired in our *Rest ^d/d^* model, such as mitogen-inducible gene 6 (*Mig6*) and matrix metalloproteinase-11 (*Mmp11*), suggesting that in the absence of REST, impaired regulation of P4 response genes is selective, and not a global effect. These findings indicate that loss of uterine REST in our mouse model is associated with impaired regulation of select P4 responsive genes, similar to gene expression documented in the endometrium of patients diagnosed with endometriosis.

**Figure 5.**
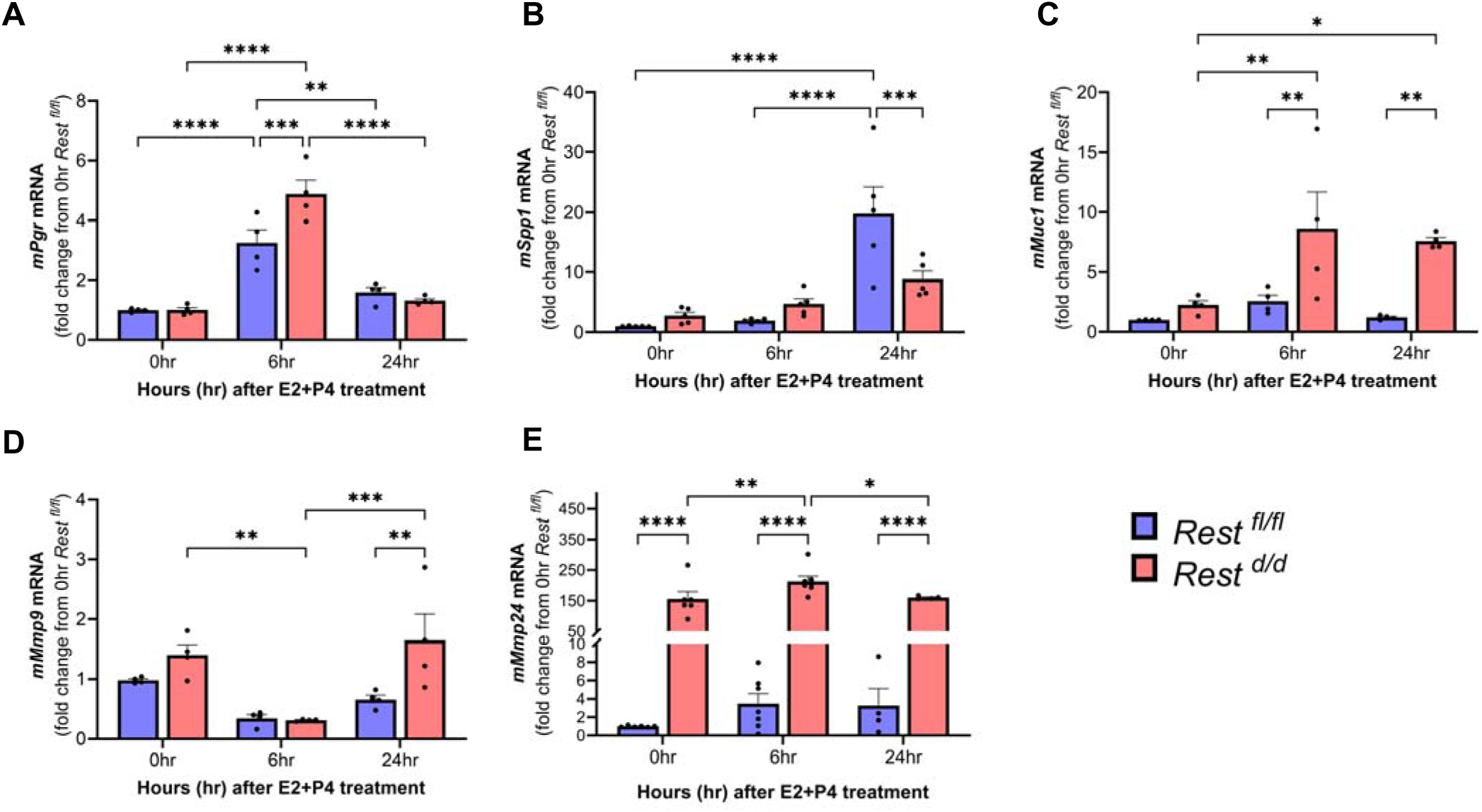
Loss of uterine REST is associated with impaired regulation of select P4 responsive genes after steroid treatment. Whole uterine tissue from *Rest ^fl/fl^* and *Rest ^d/d^* female mice were assessed for mRNA expression of P4 responsive genes (**A**) Progesterone receptor, *Pgr* (n = 4/genotype/time point) (**B**) secreted phosphoprotein-1 or osteopontin, *Spp1* (n = 5/genotype/time point) (**C**) mucin-1, *Muc1* (n = 4/genotype/time point) (**D**) matrix metalloproteinase-9, *Mmp9* (n = 4/genotype/time point) and (**E**) matrix metalloproteinase-24, *Mmp24* (0hr, n = 6/genotype; 6hrs, n = 7/genotype; 24hrs, n = 4/genotype) at 0hr, 6hrs, and 24 hrs after E2+P4 combination steroid injection. *Rest ^d/d^* fold change values normalized to average fold change from 0hr *Rest ^fl/fl^*for each gene. Uterine mRNA expressions of select P4 responsive genes are differentially regulated in the presence of E2+P4 in conditional knockout mice as compared to controls. Data was analyzed between genotype within time point and within genotype across time points using two-way ANOVA (Tukey’s multiple comparison test). *Significance is set at *p < 0.05; **p <0.01; ***p < 0.001; ****p < 0.0001*. Data is represented as mean ± SEM.

### Endometriosis-induction with Rest deficient tissue impact on pain-like behavior

Endometriosis is characterized not only by fertility challenges, but also chronic pelvic pain and heightened vaginal sensitivity. In addition to its role in tumor suppression, REST functions as a neuronal gene repressor in non-neuronal tissues, and its dysregulation can influence inflammation and pro-nociceptive environments.^39–42^ To investigate how REST deficiency in endometriosis influences pain-like behavior, we used an established protocol to induce endometriosis-like pathology in a mouse model (**Figure 6A**).^43,44^ Endometrial fragments from donor mice with *Rest ^fl/fl^* or *Rest ^d/d^* genotypic backgrounds were injected into the peritoneal cavity of wild type C57BL/6 female recipient mice where tissue would develop into endometriotic-like lesions. **Figure 6** shows the experimental design and results assessing the impact of REST deficient endometrial tissue on pain-like behavior in an endometriosis-induced mouse model, including the surgery/induction process (**Figure 6A and Methods 2.8: *Endometriosis-induction surgical procedure***), resting and treatment timeline (**Figure 6B**), hindpaw mechanical sensitivity assessment (**Figure 6C**), and vaginal sensitivity using vaginal balloon distension and measuring visceromotor response (**Figure 6D-E**). After endometriosis-induction, mice recovered from surgery and began a regime of daily placebo (sesame oil control) or P4 injections for 4 consecutive days (**Figure 6B**). On day 14, mechanical allodynia and widespread sensitivity was evaluated using hindpaw Von Frey methodology, where a lower withdrawal threshold would indicate increased sensitivity.^45–47^ In recipients of *Rest ^fl/fl^* endometrial tissue, P4 treatment significantly increased withdrawal threshold relative to recipients of *Rest ^fl/fl^*tissue treated with placebo, indicating a higher tolerance for mechanical pressure and lower mechanical sensitivity (**Figure 6C**). In contrast, *Rest ^d/d^*recipients show a moderate trend of increased threshold when treated with P4 but overall was not significant between the placebo and P4 treated groups *(p = 0.0864)*, suggesting a resistance to the effects of P4 treatment for mechanical allodynia in association with endometriosis. Recipients of *Rest ^d/d^*tissue exhibited lower withdrawal thresholds when compared to *Rest ^fl/fl^*of the same treatment. This was observed under both treatment conditions, placebo and P4 injections, indicating heightened mechanical sensitivity. On day 15, vaginal hyperalgesia was assessed using graded vaginal balloon distensions (VBD) to measure the visceromotor reflex (VMR), a validated indicator of endometriosis-associated vaginal sensitivity in rodents.^43,48,49^ In our model, no difference was observed in visceromotor reflex, whether between treatments and/or genotype of donor tissue, overall (**Figure 6D**) or individual pressures of balloon distension (**Figure 6E**), with uterine REST deficient tissue and P4 treatment having minimal effect on displays of vaginal sensitivity in our mouse model of endometriosis.

**Figure 6.**
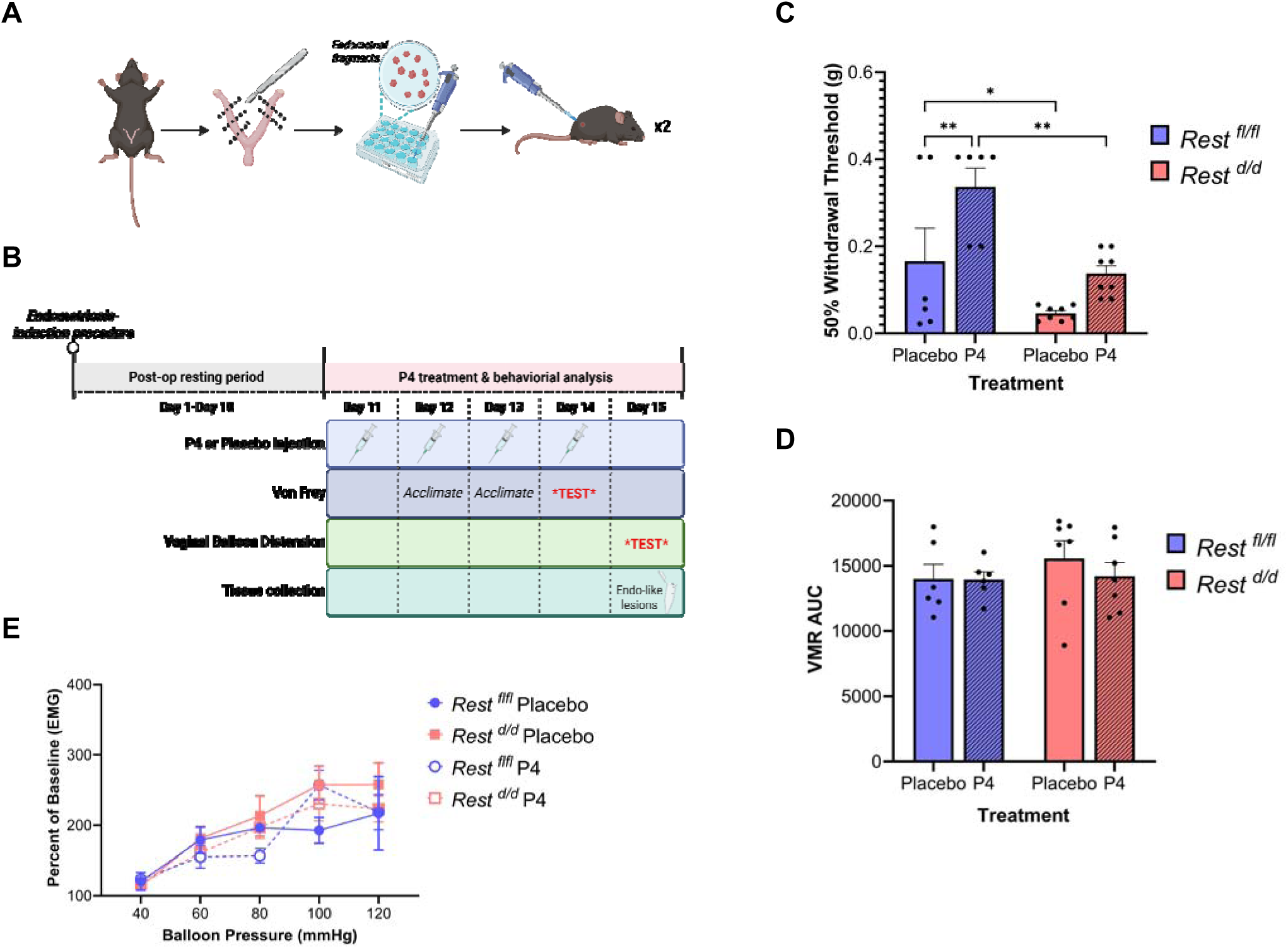
Impact of *Rest* deficient endometriotic-like lesions on pain-like behavior in an endometriosis-induced experimental mouse model. (**A**) Schematic detailing endometriosis-induction mouse model. Wild type C57BL/6 recipient female mice (3-4 months of age) were injected with endometrial tissue harvested from donor mice with genotypic background *Rest ^fl/fl^* control or *Rest ^d/d^*conditional knockout to induce endometriosis (one donor mouse produced enough fragments for two recipient mice). (**B**) Depiction of the experimental timeline and design including post-induction operation resting period, administration of placebo or P4 treatment, and pain-like behavior testing (“TEST”). (**C**) Hindpaw Von Frey (VF) behavior assessment was conducted to observe withdrawal responses using the Up-Down method (Control, n = 6/treatment; Knockout, n = 8/treatment). VF data was analyzed using two-way ANOVA followed by Fisher’s LSD test. Data are displayed as mean ± standard error of the mean (SEM) and significance set at *\*p < 0.05; **p < 0.01*. *Rest* deficient endometriotic-like lesions had a significant impact on hindpaw mechanical withdrawal threshold, with significantly lower withdrawal threshold in the mice injected with *Rest ^d/d^*endometrial tissue as compared to *Rest ^fl/fl^* control endometrial tissue in both the placebo (sesame oil) and P4 groups. (**D-E**) The visceromotor response (VMR) was measured by recording EMG activity of the abdominal musculature during graded balloon distension of the vagina (VBD) (Control, n = 6/treatment; Knockout, n = 7/treatment). Balloon distensions were administered in triplicates for the following pressures of 40, 60, 80, 100, and 120mmHg. VMR is quantified using (**D**) the area under the curve and (**E**) as a percentage of baseline EMG recording at each pressure/group. VMR data were analyzed using two-way ANOVA (with repeated measures) followed by Bonferroni’s test. Data are displayed as mean ± standard error of the mean (SEM) and significance set at alpha <0.05. No significant difference in vaginal sensitivity was observed between *Rest ^fl/fl^* and *Rest ^d/d^*recipient mice (between genotype within same treatment or between different treatments within genotype) for AUC and each individual pressure *(p > 0.05)*.

### Loss of uterine Rest results in significantly larger endometriotic-like lesions

Survival and growth of endometriotic lesions are supported by aberrant expression of P4 responsive genes and uninhibited E2 activity. However, it is well established that pain severity experienced by patients with endometriosis does not correlate to diseaseseverity.^22,23^ All mice subjected to vaginal balloon distensions were euthanized and endometriotic-like lesions were excised from surrounding tissue in the abdominal cavity. Lesion size was recorded (mm^3^) with representative images in **Figure 7A**.

**Figure 7.**
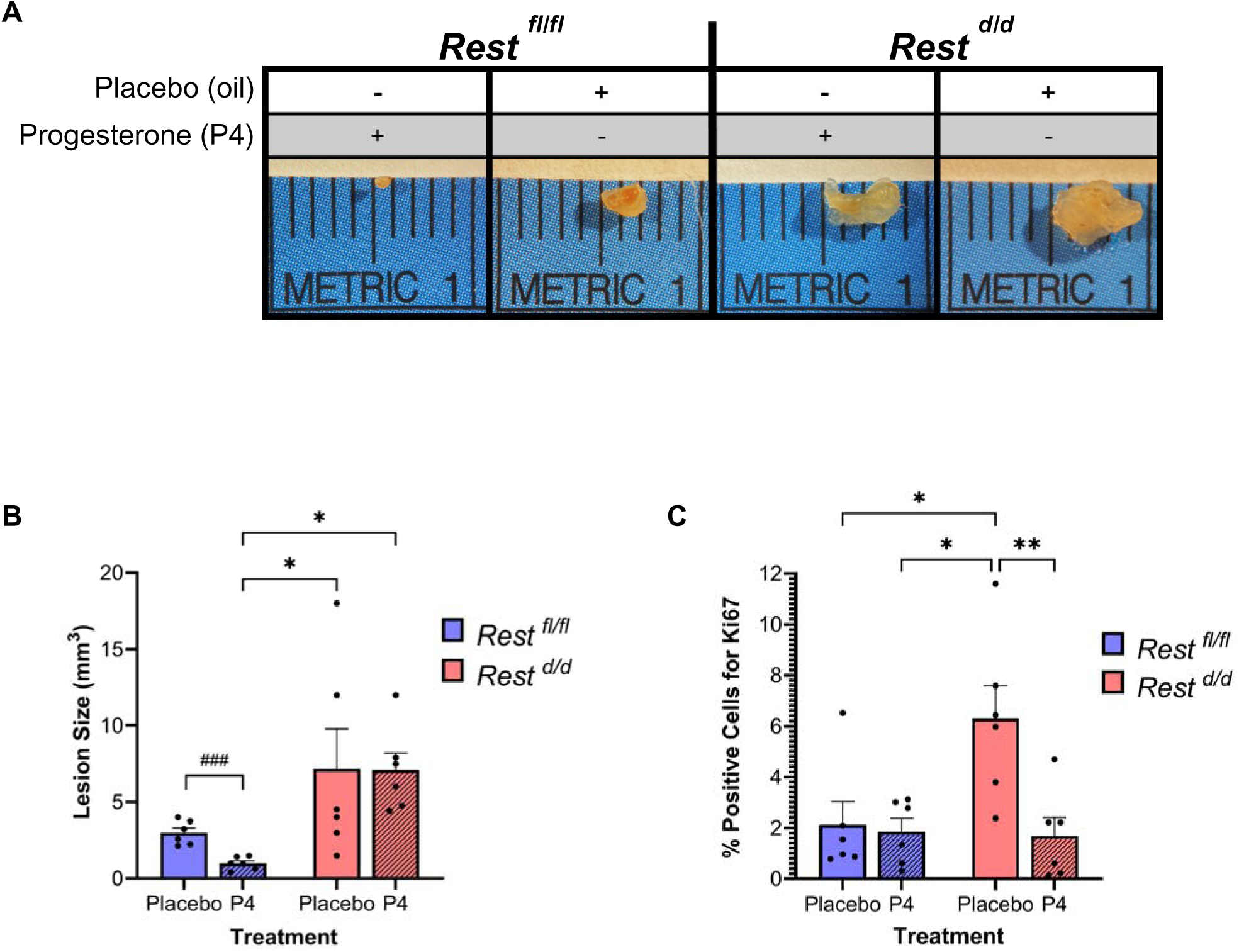
The impact of uterine tissue Rest expression and progesterone treatment in endometriotic-like lesion development. (**A**) Representative image of wild-type recipient mouse endometriotic-like lesions (*Rest ^fl/fl^* or *Rest ^d/d^*fragments) collected from the abdominal region and measured (length, width, height). (**B**) Area (mm^3^) was analyzed using Welch’s t-test between placebo and P4 treatments within the same endometrial fragment genotype *(^###^p < 0.001)*, two-way ANOVA (Tukey’s multiple comparison test) was used to analyze area (mm^3^) across all groups *(*p < 0.05; **p < 0.01)*. *Rest ^fl/fl^*, but not *Rest ^d/d^*, lesions responded to P4 treatment and significantly decreased in size as compared to placebo (###), with *Rest ^d/d^* placebo and P4 treated lesions remaining on average similar in size. Placebo and P4 treated *Rest ^d/d^* lesions remained significantly larger than P4 treated *Rest ^fl/fl^* lesions (*) (n = 6/genotype/treatment). (**C**) Immunohistochemical staining with Ki67 shows a significantly higher percentage of Ki67-positive cells in the *Rest ^d/d^* placebo group as compared to *Rest ^fl/fl^* placebo and P4 groups. Ki67-positive cells were analyzed using ImageJ followed by two-way ANOVA (Tukey’s multiple comparison test) across all groups *(*p < 0.05; **p < 0.01)* within the entire tissue cross-section of the lesion (n = 6/genotype/treatment).

*Rest ^fl/fl^* endometrial tissue lesions treated with P4 were significantly smaller relative to placebo (average 1mm^3^ vs. of 3mm^3^), unlike *Rest ^d/d^* endometrial tissue lesions which did not significantly differ in size between placebo and P4 treated groups (average 7.16mm^3^ *Rest ^fl/fl^* and 7.09mm^3^ *Rest ^d/d^*) (**Figure 7B**). Placebo treated *Rest ^d/d^* lesions exhibited a significantly higher percentage of Ki67-positive cells compared to all other groups, indicating increased cellular proliferation in these endometriotic-like lesions (**Figure 7C**). To assess the impact of REST deficiency on sex steroid receptor expression, lesions were stained for ER-alpha (**Figure 8A-D representative images**) and PGR protein (**Figure 8G-J representative images**).

**Figure 8.**
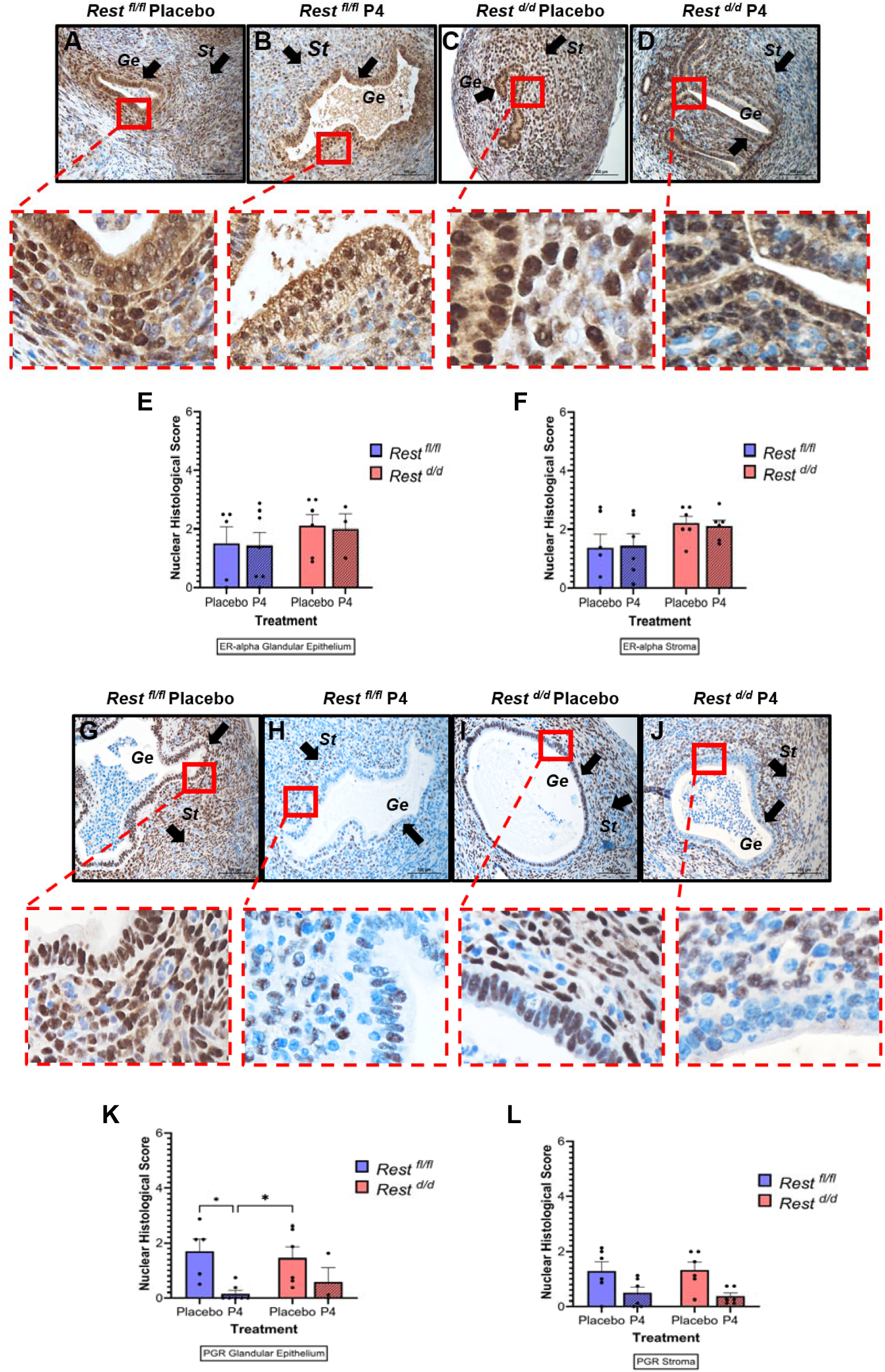
The impact of uterine tissue Rest expression and progesterone treatment in endometriotic-like lesion steroid receptor protein expression. (**A-D, G-J**) Representative immunohistochemical staining images for estrogen receptor alpha (ER-alpha) (**A-D**) and progesterone receptor (PGR) (**G-J**) in cross-sections of endometriotic-like lesions for (**A, G**) *Rest ^fl/fl^*lesion tissue from mice treated with placebo, (**B, H**) *Rest ^fl/fl^*lesion tissue from mice treated with P4, (**C, I**) *Rest ^d/d^*lesion tissue from mice treated with placebo, and (**D, J**) *Rest ^d/d^*lesion tissue from mice treated with P4 (ER-alpha/PGR, brown staining; hematoxylin, purple counterstain). Images were taken at 20X (scale bar indicates 100μm). (**E-F, K-L**) Tissue samples were assessed using histological scoring (H-score) for nuclear ER-alpha and PGR staining in endometrial stroma (St) and glandular epithelium (Ge) (black arrows) across all groups. Samples were scored on a scale of 0-3 based on previously established parameters for histological scores with 0 being little/absent staining and 3 being the most strong/intense staining. H-scores were analyzed using two-way ANOVA (Tukey’s multiple comparison test) across all groups. *Significance is set at *p < 0.05; **p <0.01; ***p < 0.001; **** p < 0.0001)*. Data is represented as mean ± SEM. *Glandular epithelium ER-alpha/PGR staining: Rest ^fl/fl^ Placebo, n = 5; Rest ^fl/fl^ P4, n = 7; Rest ^d/d^ Placebo, n = 6; Rest ^d/d^ P4, n = 3. Stromal ER-alpha/PGR staining: n = 6/genotype/treatment*.

Genotypic background of fragments and/or treatment had no significant effect on nuclear expression of ER-alpha in the glandular epithelium or stroma of the endometriotic-like lesions across all groups (**Figure 8E-F**). *Rest ^fl/fl^* P4-treated mice expressed significantly lower nuclear PGR protein in the glandular epithelium compared to placebo treated counterpart, whereas the *Rest ^d/d^* P4 group showed a trend towards lower nuclear PGR in the glandular epithelium compared to the *Rest ^d/d^*placebo treated lesions but was overall non-significant (*p = 0.4382*) (**Figure 8K**). A similar observation was made for nuclear PGR protein staining in the lesion stroma, with a non-significant trend towards lower PGR expression in the P4 treated groups compared to placebo groups in both *Rest ^fl/fl^* and *Rest ^d/d^* tissue (*p = 0.1436* and *p = 0.0580*, respectively) (**Figure 8L**).

Interpretation of these findings suggests Rest-deficient endometriotic-like lesions are receptive to P4 treatment with a response in PGR expression similar to *Rest ^fl/fl^* lesion tissue, however it is less effective due to large lesion size and increased proliferative activity observed in the *Rest ^d/d^* lesions. These results further support REST as a key regulator of P4 signaling and a potential mechanism underlying endometriosis pathology.

## Discussion

Endometriosis is a chronic estrogen-dependent disease characterized by aberrant steroid hormone signaling, particularly impaired progesterone (P4) responsiveness, which contributes to lesion development and persistence. Although P4-based therapies are widely used clinically, many patients exhibit incomplete or variable responses, highlighting a limited understanding of the molecular mechanisms underlying P4 resistance in endometriosis. Previous studies have demonstrated dysregulation of P4 target genes in the endometrium of women with endometriosis.^50^ Altered PGR isoform ratios, inflammation, and epigenetic mechanisms have all been proposed to disrupt P4 signaling, although the underlying molecular pathways remain unclear.^12^ Infertility and chronic pain are among the most common symptoms reported by patients, yet their presentation varies. This variability may reflect differences in hormone responsiveness that influence disease presentation, progression, and treatment response.

RE-1 silencing transcription factor (REST) has emerged as a candidate regulator of steroid hormone signaling through its interaction with PGR and co-regulation of P4-responsive genes. Given that reduced REST expression has been reported in uterine leiomyoma and breast cancer, we hypothesized that REST is similarly reduced in endometriosis and contributes to impaired P4 responsiveness and lesion persistence. To test this, we characterized REST expression in human endometriosis tissues and used a uterine-specific Rest knockout mouse model to investigate its role in fertility, hormone responsiveness in the uterine tissue, endometriotic-like lesion development, and pain-related phenotypes.

Our study identifies, for the first time, loss of functional nuclear REST protein as a feature of both eutopic endometrium and ectopic endometriotic lesions when compared to healthy control endometrium. In human endometriosis tissues, REST protein was markedly reduced in the nucleus and mislocalized to the cytoplasm in both eutopic and ectopic tissue. Cytoplasmic retention of REST may contribute to this functional loss by promoting ubiquitination-mediated degradation.^51^ These findings support REST as a potential regulator of P4 responsiveness in endometriosis. However, elucidating the mechanisms underlying REST protein loss in endometriosis will also be important. Targeting these processes may provide new therapeutic opportunities to restore REST function.

Using our conditional uterine Rest knockout, loss of REST resulted in progressive subfertility, supporting a role for REST in reproductive function. Although Rest knockout females retained partial fertility during early breeding cycles, reproductive success declined over time compared with control females, which maintained consistently high fertility across all cycles. This pattern may reflect transient compensatory mechanisms or delayed Cre-mediated deletion. Because PGR is also expressed in the ovary, we cannot exclude potential ovarian contributions, including altered oocyte production. Nevertheless, the phenotype is consistent with the subfertility frequently associated with endometriosis and supports the utility of this model for studying endometriosis-associated reproductive dysfunction.

At the tissue level, Rest deficiency produced a hyper-estrogenic uterine phenotype and reduced P4-mediated suppression of estrogen-driven uterine growth, observed during both chronic and combined hormone treatment. Importantly, this phenotype does not suggest complete P4 resistance, but rather a blunted or diminished responsiveness to P4. This distinction is critical, as it more closely resembles the partial P4 insensitivity observed clinically in endometriosis. Our study is a step forward in identifying pathways involved in the characterization of endometriosis associated subfertility, which can lead to better diagnostic measures for fertility outcomes in endometriosis patients.

Consistent with studies showing dysregulation of P4 target genes in women with endometriosis,^50^ uterine Rest loss selectively altered expression of P4-responsive genes in mouse tissue. This selective effect suggests that REST functions as a context-dependent transcriptional regulator, likely through interaction with PGR at shared regulatory elements. Notably, impaired P4 responsiveness occurred despite preserved PGR expression, indicating that P4 resistance can develop independently of receptor abundance. Loss of REST did not significantly alter PGR transcript levels, further supporting a role for REST in epigenetic or co-regulatory control of P4 target gene expression. These findings are consistent with previous reports demonstrating blunted P4 signaling responsiveness despite minimal differences in PGR expression between eutopic endometrium from women with and without endometriosis.^20^

Our lesion data further supports this interpretation. Estrogen receptor-alpha and P4 receptor expression were unchanged in endometriotic-like lesions derived from Rest-deficient tissue. However, lesion size and proliferation remained largely unaffected by P4 treatment compared with lesions from control mice. Thus, endometriotic-like tissue receptor expression alone in this model was insufficient to drive an effective biological response to P4 treatment in the absence of REST. This suggests that REST may facilitate proper PGR transcriptional activity, potentially through chromatin remodeling or co-regulatory complex formation, and influence tissue P4 responsiveness. Similar observations have been reported in other hormone-dependent systems, such as breast cancer models, where steroid receptor expression remains stable despite altered downstream signaling.^26–28,34^

Pain-related outcomes only partially correlated with lesion characteristics. Mechanical sensitivity increased in mice receiving Rest-deficient endometrial tissue, whereas vaginal sensitivity remained unchanged. This dissociation is consistent with clinical observations that lesion burden does not necessarily correlate with pain severity and suggests that REST may differentially influence peripheral and visceral pain pathways. Variability in P4 responsiveness associated with REST expression may also contribute to differences in pain phenotypes. Additional studies incorporating inflammatory analyses alongside REST expression and pain assessments will be important to clarify these relationships.

Collectively, these findings support a model in which loss of uterine Rest results in a blunted P4 signaling responsiveness, persistent estrogenic activity, and enhanced lesion survival. This framework also raises the possibility that REST contributes to the heterogeneity observed in endometriosis symptoms and treatment responses.

Considerations when interpreting the results from this study include variability in human endometriotic lesion type, anatomical location, and duration each lesion has been present may contribute to differences in REST expression and were not fully stratified. Although the conditional knockout model provides important mechanistic insight, it does not fully recapitulate the complexity of human endometriosis, including systemic immune influences, environmental factors, and lesion heterogeneity. Future studies should include broader characterization of P4-responsive genes within REST-dependent transcriptional networks beyond the scope of this work. In addition, some endometriotic-like lesion cross-sections lacked well-defined glandular structures, which may affect interpretation of immunohistochemical analyses. Lesions were evaluated at a single time point (two weeks following experimental induction), limiting assessment across different stages of lesion establishment and progression.

Assessment of pain-like behavior also revealed limited and modality-specific effects. The absence of changes in vaginal sensitivity may reflect model limitations or suggest that REST primarily influences specific nociceptive pathways. Clinically, these findings raise the possibility that REST status could serve as a biomarker for P4 responsiveness or treatment stratification. If REST contributes to variability in hormone responsiveness, it may help explain why some patients respond poorly to P4-based therapies and endometriosis present with a variation in fertility and pain symptomology. Integrating REST into broader models of endometriosis pathophysiology, including immune, inflammatory, and neurogenic pathways, will also be important for understanding symptom heterogeneity. Overall, our findings support a model in which loss of REST promotes blunted P4 responsiveness, persistent estrogenic activity, and enhanced lesion persistence, key features underlying endometriosis pathophysiology and clinical variability. More advanced translational models may help determine whether restoring REST function can rescue P4 responsiveness and limit disease progression.

## Materials and Methods

### Human subjects and tissue acquisition

All use of human tissues was performed in accordance with Institutional Review Board (IRB)-approved study protocols (IRB00000161). All human endometrial tissue specimens were collected at the University of Kansas Medical Center by trained surgical professionals in the Department of Obstetrics and Gynecology. Eutopic and ectopic endometrial tissue specimens were collected from consenting women with confirmed laparoscopic endometriosis who presented with pelvic pain following failed previous endometriosis treatment and undergoing surgical removal of endometriotic lesion tissue. Control eutopic endometrial tissue specimens were collected from women with pain but no laparoscopic evidence of endometriosis. For some experiments, archived endometrial and endometriotic lesions tissue blocks were retrieved from the Department of Pathology and Laboratory Medicine for immunohistochemical localization studies. Patients who reported the use of hormones and/or birth control were excluded. Information on stage of menstrual cycle (Proliferative or Secretory) was based on patient’s medical records and histological assessment. Grouping was done on basis of stage of menstrual cycle (Proliferative vs. Secretory). Additional patient characteristics were noted including age, presence of leiomyoma, adenomyosis, and/or polyps, and stage of endometriosis (Stage I/II vs. Stage II/IV) as identified by pathology reports (**Table 1**).

### Human immunohistochemistry staining

Human uterine tissue specimens were processed through the Histology Core at the University of Kansas Medical Center. Formalin-fixed, paraffin-embedded tissues were sectioned at 5μm and mounted on Fisherbrand Superfrost Plus microscope slides (Thermo Fisher Scientific, Waltham, MA, cat#12-550-15). Tissue slides were deparaffinized and rehydrated using xylene, 100/95/70% ethanol, and tap water. Slides were then incubated in antigen retrevial (Antigen Unmasking Solution, Vector Laboratories, cat#H-3300) at 95C for 10 minutes. Tissue was subjected to immunohistochemical (IHC) protein localization for REST (Proteintech, Rosemont, IL, cat#22242-1-AP; 1:200 dilution). IHC was performed following the recommendations of the manufacturer using VectaStain ABC system (Vector laboratories, Newark, CA, cat#PK-6106) and ImmPACT DAB Substrate Kit (Vector Laboratories, cat#SK-4105). Slides were counterstained with Hematoxylin (Vector Laboratories, cat#H-3404) and cover-slipped using non-aqueous Vectamount mounting media (Vector Laboratories, cat#H-5000).

### Histological scoring (H-Score) for human tissue

Protein localization was identified as a positive reaction by a dark brown coloring on the tissue slides against purple Hematoxylin counterstain. Antibody intensity and percent positive reaction in nuclei and cytoplasm of the endometrial glandular epithelium and stroma were quantified using the H-score method.^52^ Stained slides were assessed using a Nikon Alphaphot 2 YS2 microscope at 20X magnification. Samples were scored on a scale of 0-3, in increments of 0.25, based on previously established parameters for histological scores with 0 being little/absent staining and 3 being the most strong/intense staining with high percent coverage. Two independent observers analyzed the stained tissue slides and values were averaged for the final H-score in both the nucleus and cytoplasm. Glands and stroma were analyzed separately for both secretory and proliferative tissue samples.

### Rest conditional knockout mouse generation and genotyping

All experiments involving animals were approved by the Unviersity of Kansas Medical Center Animal Care and Use Committee and conducted in accordance with applicable guidelines. C57BL/6 mice were housed within environmentally controlled conditions under the supervision of a licensed veterinarian following guidelines as suggested in the “Guide for the care and use of laboratory animals” by the National Research Council of the National Academies. *Rest ^fl/fl^* C57BL/6 mice were generated in the Dr. Chennathukuzhi lab. Generation of *Rest ^d/d^ (Rest ^fl/fl^ Pgr ^Cre/+^)* conditional knockout mice occured using the Cre-LoxP system and crossing *Rest ^fl/fl^* C57BL/6 mice with *Pgr ^Cre/+^*C57BL/6 mice.^53^ *Rest ^fl/fl^*mice were generated using the methods described here and in previous studies.^34,54,55^ Exon 2 is deleted from the *Rest* gene in mice homozygous for *Rest* (*Rest ^fl/fl^)* and heterozygous for progesterone receptor Cre knock-in (*Pgr ^Cre/+^*), creating a truncated non-functional form of Rest in tissue where the progesterone receptor Cre is expressed. Ear punches were taken and the mice were genotyped by PCR for *Rest* using forward primer (5’-TGTAGTTTCCAAACTGTGACTTCG -3’) and two reverse primers (Reverse1: 5’-TGAACTGATGGCGAGCTCAGACC-3’) (Reverse2: 5’-GCTACAAAATGCGTAAGTTCAAGG-3’). DNA was extracted from ear punches using REDExract-N-Amp kit (Sigma-Aldrich, St.Louis, MO, cat#XNAT-1KT) and PCR was performed according to manufacture’s instructions. The PCR amplification cycle was run as follows: Initial denaturation for 95°C for 5 mins and the next steps are repeated 35 times which are denaturization for 95°C for 30 secs followed by annealing and extension at 60°C and 72°C respectively for 30 secs each. After completing 35 cycles, final extension is at 72°C for 10 mins followed by hold at 4°C. Samples, ladder, and controls were then run on 1% agarose gel at 80 volts for half an hour and visualized using BioRad GelDoc (Bio-Rad Laboratories, Hercules, CA).

### Female conditional knockout mouse fertility trials

Mature 2-month-old *Rest ^d/d^ (Rest ^fl/fl^ Pgr ^Cre/+^)* female mice were mated with wild type C57BL/6 male of proven fertility to assess ability of *Rest^d/d^* to produce offspring. *Rest ^fl/fl^* control female mice were mated with C57BL/6 males as a control for comparison. Females of both genotypes were checked daily for presence of vaginal plug indicative of successful mating. Females who displayed vaginal plugs were separated from breeding cages. Once separated, mice were checked daily, and the date of delivery and number of living pups were recorded. Pups were weaned after 21 days, and female dams were remated. After 25 days, females that failed to deliver pups were marked as “no litter” and remated. Females underwent 3 consecutive breeding cycles over a course of 4 months. The age for females after the third breeding cycle did not exceed 6 months.

### Subcutaneous steroid injection treatments

#### Early and late effects of steroid treatment

Fourteen days after ovariectomy, mice were injected subcutaneously with estradiol-17β (E2, 10 μg/kg BW; Sigma-Aldrich, St.Louis, MO, cat#E8875-1G) or E2 + progesterone/P4 (P4, 100 mg/kg BW; Sigma-Aldrich, St.Louis, MO, cat#P8737-5G) (E2, 10 μg/kg BW + P4, 100 mg/kg BW). Mice were then euthanized at 6hrs or 24hrs after injection to evaluate early and late effects of steroid treatments. Vehicle injections (0.1 ml sesame oil) were administered for 0hr time points as a control/baseline. Uteri were removed, trimmed of fat and connective tissue, weighed, and then stored in RNAlater solution (Invitrogen/Thermo Fisher, Waltham, MA, cat#AM7024) at 20°C until whole uterine horn tissue was processed for total RNA extraction.

#### Chronic progesterone treatment experiments

To assess the impact of chronic P4 treatment, separate groups of mice were injected with vehicle (0.1 ml sesame oil; Sigma-Aldrich, St.Louis, MO, cat#S3547) or P4 (100 mg/kg BW) three consecutive times, 24hrs apart, and euthanized 24hrs after the last injection. Uteri were removed, trimmed of fat and connective tissue, weighed, and then stored in RNAlater solution at 20°C until whole uterine horn tissue was processed for total RNA extraction.

### Mouse uterine horn RNA isolation and RT-qPCR

For assessment of mRNA expression, total RNA was isolated from whole uterine horn tissue using TRIZol reagent (Invitrogen/Thermo Fisher, Waltham, MA, cat#15596-018) and RNA pellet was resuspended in nuclease-free water. Total RNA (1ug/20uL) was used for reverse transcriptase with M-MLV Reverse Transcriptase kit (Invitrogen, cat#28025-013), Random Primers (3ug/uL, Invitrogen, cat#48190-011), and dNTP (100mM; Invitrogen/Thermo Fisher, Waltham, MA, cat#R0181). Relative gene expression of progesterone response genes was determined by qRT-PCR on a QuantStudio 7 Flex System (Thermo Fisher Scientific, Foster City, USA) using PowerSYBR Green PCR Master Mix (Applied Biosystems, Thermo Fisher Scientific, cat#4367659) and the following forward and reverse mouse primer sequences: *Pgr*, Forward (5’-CTACTCGCTGTGCCTTACCATG-3’) and Reverse (5’-CTGGCTTTGACTCCTCAGTCCT-3’); *Spp1*, Forward (5’-GCTTGGCTTATGGACTGAGGTC-3’) and Reverse (5’-CCTTAGACTCACCGCTCTTCATG-3’); *Muc1*, Forward (5’-GCCCCTTCCCGCCTGTTCAC-3’) and Reverse (5’-TCACTTGGAAGGGCAAGAAAACCTTT-3’); *Mmp9*, Forward (5’-GCTGACTACGATAAGGACGGCA-3’) and Reverse (5’-TAGTGGTGCAGGCAGAGTAGGA-3’); *Mmp24*, Forward (5’-CCTTCATCAGCAAGGAAGGATATTA-3’) and Reverse (5’-TCCAGTCACGCAGGATGTTG-3’). All samples were run in triplicate, and the average was used to calculate fold change (2^-ΔΔCT^) using the housekeeping gene *Rpl13*, Forward (5’-TACCAGAAAGTTTGCTTACCTGGG-3’) and Reverse (5’-TGCCTGTTTCCGTAACCTCAAG-3’).

### Endometriosis-induction surgical procedure^43^

Female <u>donor mice</u> (22-24 days old), of genotypic background *Rest ^fl/fl^* control or *Rest ^d/d^ (Rest ^fl/fl^ Pgr ^Cre/+^)* conditional knockout, were injected subcutaneously with 100ul of 20 IU/mL PMSG (pregnant mare’s serum gonadotropin; 2 IU/mouse; ProspecBio, East Brunswick, NJ, cat#hor-272-b) in sterile PBS to stimulate endogenous estrogen production and prime the endometrium for dissection from myometrium. 40 hours post-injection, donor mice were euthanized, and uterine horn was collected. Each uterine horn was cut into 5-6 smaller sections, and endometrial fragments (stroma and epithelium) were cut away from myometrium at the ends of each section. Collected endometrium was cut into 10 equal (5mg, 1mm^3^) fragments and resuspended in 300 µL of sterile PBS. Endometriosis was induced in 2-month-old, immune-competent, reproductively intact C57BL/6 wild-type control <u>recipient mice</u> using endometrial tissue from <u>donor mice</u>. One donor mouse produced enough endometrial tissue fragments for 2 recipient mice. Recipient females were anesthetized under isoflurane inhalation and given 100ul of 1mg/kg Meloxicam analgesic subcutaneously before surgery and again 24 hours later. A surgical area below the right ribcage was cleansed with betadine and 70% ethanol and clipped free of hair. A small incision (approximately 1.5 centimeters) was made at this location and donor endometrial tissue fragments in PBS suspension were dispersed along the peritoneal wall in the pelvic cavity via a pipettor equipped with a sterile 1000µL tip. The incision was then closed with wound clips and recipients were left to rest for 5 minutes on isoflurane inhalation before being removed from isoflurane. Mice were left in home cages for 14 days to allow endometriotic-like lesions to establish before lesion collection on day 15 post-surgery induction.

### Assessment of pain-like behavior in mice

#### Endometriosis-induction mouse groups

At 11 days post-surgery, mice began subcutaneous injections of progesterone (P4; 100 mg/kg BW) or vehicle (0.1 ml sesame oil) for four consecutive days (days 11-14 post-surgery induction), 24h apart.

#### Von Frey Filament Testing

On days 12 and 13 post-surgery induction, mice were acclimated to Von Frey conditions set up by resting them for 30 minutes each day (sitting on wire-mesh platform, white noise machine, presence of researcher who will be performing the test). Mice were placed inside individual clear plastic containers, leaving the wire-mesh exposed with access to the hindpaw. To assess referred somatic allodynia and general tenderness, mechanical pressure was applied to the hindpaw using standard Von Frey monofilaments (1.65, 2.36, 2.83, 3.22, 3.61, 4.08, 4.31, and 4.74 grams of force, Stoelting, Co., Wood Dale, IL) to the area and recording withdrawal behavior (jump, jerk, spread paw, or licking of testing area) as a positive response using the “Up-Down” method.^45,46^ After the first recorded positive response, an additional four applications of monofilaments were administered for a total of five applications. The series of responses to filament pressure were used to calculate a 50% g threshold for each mouse.^45–47^

#### Visceromotor Reflex/Vaginal Balloon Distension

On day 15 post-surgery induction, vaginal balloon distension (VBD) was administered to measure visceromotor response following a previously published protocol.^43,47,54^ While mice were under anesthesia (inhaled isoflurane), two stainless steel electrode wires were acutely implanted into the right and left lateral abdominal musculature using a 26-gauge needle and a custom-made latex balloon (1cm in length) was inserted into the vaginal canal and inflated using pressure-controlled nitrogen. Mice were placed into a Broome-style restraint (Kent Scientific, Torrington, CT) and allowed to acclimate for 30 minutes after waking from anesthesia. Balloon distensions were administered for 20 seconds in triplicate for the following pressures of 40, 60, 80, 100, and 120mmHg in 4-minute intervals. The implanted abdominal electrodes were attached to an A-M Systems differential AC Amplifier (Model 1700, A-M Systems, Sequim, WA). Electrode activity was recorded from amplified electromyographs (EMG), converted to digital reads (1401 Micro D-A converter), and then processed using Spike2 7.0 software (Cambridge Electronic Design, Cambridge, UK). VMR was quantified using the area under the curve for the 20 second distension timeframe and expressed as a percentage of baseline activity (measured 10 seconds prior to distension period).

### Endometriotic-like lesion collection and immunohistochemistry

Immediately following the completion of vaginal balloon distension/visceromotor reflex assessment, mice were euthanized, and ectopically established endometriotic-like lesion tissue was collected. Starting at the pelvis, a lower midline incision was made, and the abdominal wall and cavity were inspected for endometriotic-like lesions formed from the injected endometrial tissue near where the previous surgical incision was made. Cystic endometriotic-like lesions were excised from surrounding tissue and placed on a petri dish in sterile phosphate buffered saline (PBS) to avoid drying out while measurements were taken (area mm^3^) with a ruler before placing tissue in 10% neutralized buffered formalin. Fixed lesion tissue was paraffin-embedded and subjected to immunohistochemistry using antibodies for cellular proliferation, Ki67 (Invitrogen/Thermo Fisher, Waltham, MA, cat#MA5-14520, dilution 1:100) and localization of steroid receptors estrogen receptor-alpha (ER-□; Abcam, Cambridge, UK, cat#ab-32063, dilution 1:2000) and progesterone receptor (PGR) (Cell Signaling Technologies, Danvers, MA, cat#8757, dilution 1:500). Immunohistochemistry was performed according to the recommendations of the manufacturer using VectaStain ABC system (Vector laboratories, cat#PK-6106) and ImmPACT DAB Substrate Kit (Vector Laboratories, cat#SK-4105). Slides were counterstained with Hematoxylin (Vector Laboratories, cat#H-3404) and cover-slipped using non-aqueous Vectamount media (Vector Laboratories, cat#H-5000). Nuclear receptor protein localization was identified as dark brown coloring in the tissue against the purple hematoxylin counterstain.

### Statistical analysis

All data are presented as mean ± standard error of the mean (SEM). Statistical analyses were performed using GraphPad Prism (version 10.2.3; GraphPad Software, San Diego, CA, USA) and Microsoft Excel. Statistical significance was defined as *p* < 0.05. Significance values are indicated as follows: *\*p < 0.05; **p <0.01; ***p < 0.001; ****p < 0.0001*. All comparison analyses used two-way ANOVA followed by Tukey’s multiple comparisons (REST/PGR/ER-alpha H-Score protein assessment, Ki67, mRNA expression levels, number of pups from fertility experiments, and uterine wet weight studies). For mRNA expression analyses, all samples were run in triplicate, and average Ct values were used to calculate fold change using the 2^-ΔΔ^Ct method with *Rpl13* as the housekeeping gene. VF data was analyzed using two-way ANOVA followed by Fisher’s LSD post hoc test. Data are displayed as mean ± standard error of the mean (SEM). Visceromotor reflex (VMR) data were analyzed using two-way ANOVA (with repeated measures) followed by Bonferroni’s multiple comparisons test. All data are displayed as the mean ± standard error of the mean (SEM) and significance was set at alpha <0.05. Lesion size (area, mm^3^) was analyzed using Welch’s t-test between placebo and P4 treatments within the same endometrial fragment genotype *(^###^p < 0.001)*, two-way ANOVA (Tukey’s multiple comparison test) was used to analyze area across all groups.

## Acknowledgments

W.B.N. and V.C. were supported by grants from the NIH: WBN: R21HD099364, R01 HD105714 V.C.: P20 RR016475, R01HD113717, R01 HD094373, R01HD076450. The authors acknowledge The University of Kansas Genomics Core supported by S10 High-End Instrumentation Grant (S10 OD036343) and Frontiers CTSA grant (UL1 TR002366) at the University of Kansas Medical Center.

## References

1. Zondervan, K. T., Becker, C. M. & Missmer, S. A. Endometriosis. N Engl J Med 382, 1244–1256 (2020).

2. Agarwal, S. K. et al. Clinical diagnosis of endometriosis: a call to action. Am J Obstet Gynecol 220, 354 e1–354 e12 (2019).

3. Yoo, J. Y. et al. KRAS Activation and over-expression of SIRT1/BCL6 Contributes to the Pathogenesis of Endometriosis and Progesterone Resistance. Sci Rep 7, 6765 (2017).

4. Evans-Hoeker, E. et al. Endometrial BCL6 Overexpression in Eutopic Endometrium of Women With Endometriosis. Reprod Sci 23, 1234–41 (2016).

5. Yoo, J.-Y. et al. CRISPLD2 is a target of progesterone receptor and its expression is decreased in women with endometriosis. PLoS One 9, e100481 (2014).

6. Pabona, J. M. P. et al. Krüppel-like factor 9 and progesterone receptor coregulation of decidualizing endometrial stromal cells: implications for the pathogenesis of endometriosis. J Clin Endocrinol Metab 97, E376–392 (2012).

7. Burney, R. O. et al. Gene Expression Analysis of Endometrium Reveals Progesterone Resistance and Candidate Susceptibility Genes in Women with Endometriosis. Endocrinology 148, 3814–3826 (2007).

8. Reis, F. M., Coutinho, L. M., Vannuccini, S., Luisi, S. & Petraglia, F. Is Stress a Cause or a Consequence of Endometriosis? Reprod Sci 27, 39–45 (2020).

9. Bedaiwy, M. A. et al. Abundance and Localization of Progesterone Receptor Isoforms in Endometrium in Women With and Without Endometriosis and in Peritoneal and Ovarian Endometriotic Implants. Reprod Sci 22, 1153–1161 (2015).

10. Bulun, S. E. et al. Molecular biology of endometriosis: from aromatase to genomic abnormalities. Semin Reprod Med 33, 220–224 (2015).

11. Attia, G. R. et al. Progesterone receptor isoform A but not B is expressed in endometriosis. J Clin Endocrinol Metab 85, 2897–2902 (2000).

12. Patel, B. G., Rudnicki, M., Yu, J., Shu, Y. & Taylor, R. N. Progesterone resistance in endometriosis: origins, consequences and interventions. Acta Obstet Gynecol Scand 96, 623–632 (2017).

13. Lessey, B. A. & Kim, J. J. Endometrial receptivity in the eutopic endometrium of women with endometriosis: it is affected, and let me show you why. Fertil Steril 108, 19–27 (2017).

14. Nothnick, W. & Alali, Z. Recent advances in the understanding of endometriosis: the role of inflammatory mediators in disease pathogenesis and treatment. F1000Res 5, F1000 Faculty Rev-186 (2016).

15. Borghese, B., Zondervan, K. T., Abrao, M. S., Chapron, C. & Vaiman, D. Recent insights on the genetics and epigenetics of endometriosis. Clin Genet 91, 254–264 (2017).

16. Giudice, L. C. & Kao, L. C. Endometriosis. Lancet 364, 1789–99 (2004).

17. Burney, R. O. & Giudice, L. C. Pathogenesis and pathophysiology of endometriosis. Fertil Steril 98, 511–9 (2012).

18. Tanbo, T. & Fedorcsak, P. Endometriosis-associated infertility: aspects of pathophysiological mechanisms and treatment options. Acta Obstet Gynecol Scand 96, 659–667 (2017).

19. Donnez, J., Donnez, O., Orellana, R., Binda, M. M. & Dolmans, M. M. Endometriosis and infertility. Panminerva Med 58, 143–150 (2016).

20. Da Broi, M. G. & Navarro, P. A. Oxidative stress and oocyte quality: ethiopathogenic mechanisms of minimal/mild endometriosis-related infertility. Cell Tissue Res 364, 1–7 (2016).

21. Lessey, B. A., Lebovic, D. I. & Taylor, R. N. Eutopic endometrium in women with endometriosis: ground zero for the study of implantation defects. Semin Reprod Med 31, 109–124 (2013).

22. Shakiba, K., Bena, J. F., McGill, K. M., Minger, J. & Falcone, T. Surgical treatment of endometriosis: a 7-year follow-up on the requirement for further surgery. Obstet Gynecol 111, 1285–92 (2008).

23. Li, T. et al. Endometriosis alters brain electrophysiology, gene expression and increases pain sensitization, anxiety, and depression in female mice. Biol Reprod 99, 349–359 (2018).

24. Greaves, E. et al. Estradiol is a critical mediator of macrophage-nerve cross talk in peritoneal endometriosis. Am J Pathol 185, 2286–97 (2015).

25. Yilmaz, B. D. & Bulun, S. E. Endometriosis and nuclear receptors. Hum Reprod Update 25, 473–485 (2019).

26. Cloud, A. S. et al. Loss of REST in breast cancer promotes tumor progression through estrogen sensitization, MMP24 and CEMIP overexpression. BMC Cancer 22, 180 (2022).

27. Cloud, A. S. et al. Loss of the repressor REST affects progesterone receptor function and promotes uterine leiomyoma pathogenesis. Proc. Natl. Acad. Sci. U.S.A. 119, e2205524119 (2022).

28. Nothnick, W. B. & Chennathukuzhi, V. The molecular role of RE1 silencing transcription factor in uterine diseases: is there potential for targeted therapeutic development? Expert Opinion on Therapeutic Targets 29, 605–607 (2025).

29. Seth, K. A. & Majzoub, J. A. Repressor element silencing transcription factor/neuron-restrictive silencing factor (REST/NRSF) can act as an enhancer as well as a repressor of corticotropin-releasing hormone gene transcription. J Biol Chem 276, 13917–13923 (2001).

30. Abramovitz, L. et al. Dual role of NRSF/REST in activation and repression of the glucocorticoid response. J Biol Chem 283, 110–119 (2008).

31. Ballas, N., Grunseich, C., Lu, D. D., Speh, J. C. & Mandel, G. REST and its corepressors mediate plasticity of neuronal gene chromatin throughout neurogenesis. Cell 121, 645–657 (2005).

32. Garcia-Manteiga, J. M., D’Alessandro, R. & Meldolesi, J. News about the Role of the Transcription Factor REST in Neurons: From Physiology to Pathology. Int J Mol Sci 21, (2019).

33. Kaya, H. S. et al. Roles of progesterone receptor A and B isoforms during human endometrial decidualization. Mol Endocrinol 29, 882–895 (2015).

34. Varghese, B. V. et al. Loss of the repressor REST in uterine fibroids promotes aberrant G protein-coupled receptor 10 expression and activates mammalian target of rapamycin pathway. Proc Natl Acad Sci U S A 110, 2187–92 (2013).

35. Shimojo, M. & Hersh, L. B. REST/NRSF-interacting LIM domain protein, a putative nuclear translocation receptor. Mol Cell Biol 23, 9025–9031 (2003).

36. Shimojo, M. RE1-silencing transcription factor (REST) and REST-interacting LIM domain protein (RILP) affect P19CL6 differentiation. Genes Cells 16, 90–100 (2011).

37. Graham, A., Falcone, T. & Nothnick, W. B. The expression of microRNA-451 in human endometriotic lesions is inversely related to that of macrophage migration inhibitory factor (MIF) and regulates MIF expression and modulation of epithelial cell survival. Hum Reprod 30, 642–652 (2015).

38. Krikun, G. et al. A novel immortalized human endometrial stromal cell line with normal progestational response. Endocrinology 145, 2291–2296 (2004).

39. Reddy, B. Y., Greco, S. J., Patel, P. S., Trzaska, K. A. & Rameshwar, P. RE-1-silencing transcription factor shows tumor-suppressor functions and negatively regulates the oncogenic TAC1 in breast cancer cells. Proc Natl Acad Sci U S A 106, 4408–13 (2009).

40. Uchida, H., Ma, L. & Ueda, H. Epigenetic gene silencing underlies C-fiber dysfunctions in neuropathic pain. J Neurosci 30, 4806–14 (2010).

41. Zhang, J., Chen, S. R., Chen, H. & Pan, H. L. RE1-silencing transcription factor controls the acute-to-chronic neuropathic pain transition and Chrm2 receptor gene expression in primary sensory neurons. J Biol Chem 293, 19078–19091 (2018).

42. Su, X. J. et al. Roles of the Neuron-Restrictive Silencer Factor in the Pathophysiological Process of the Central Nervous System. Front Cell Dev Biol 10, 834620 (2022).

43. Alali, Z. et al. 60S acidic ribosomal protein P1 (RPLP1) is elevated in human endometriotic tissue and in a murine model of endometriosis and is essential for endometriotic epithelial cell survival in vitro. Mol Hum Reprod 26, 53–64 (2020).

44. Nothnick, W. B., Graham, A., Holbert, J. & Weiss, M. J. miR-451 deficiency is associated with altered endometrial fibrinogen alpha chain expression and reduced endometriotic implant establishment in an experimental mouse model. PLoS One 9, e100336 (2014).

45. Chaplan, S. R., Bach, F. W., Pogrel, J. W., Chung, J. M. & Yaksh, T. L. Quantitative assessment of tactile allodynia in the rat paw. J Neurosci Methods 53, 55–63 (1994).

46. Dixon, W. J. Efficient analysis of experimental observations. Annu Rev Pharmacol Toxicol 20, 441–62 (1980).

47. Pierce, A. N., Ryals, J. M., Wang, R. & Christianson, J. A. Vaginal hypersensitivity and hypothalamic-pituitary-adrenal axis dysfunction as a result of neonatal maternal separation in female mice. Neuroscience 263, 216–30 (2014).

48. Berkley, K. J., Cason, A., Jacobs, H., Bradshaw, H. & Wood, E. Vaginal hyperalgesia in a rat model of endometriosis. Neurosci Lett 306, 185–8 (2001).

49. McAllister, S. L., McGinty, K. A., Resuehr, D. & Berkley, K. J. Endometriosis-induced vaginal hyperalgesia in the rat: role of the ectopic growths and their innervation. Pain 147, 255–64 (2009).

50. Burney, R. O. et al. Gene expression analysis of endometrium reveals progesterone resistance and candidate susceptibility genes in women with endometriosis. Endocrinology 148, 3814–3826 (2007).

51. Huang, Z. & Bao, S. Ubiquitination and Deubiquitination of REST and Its Roles in Cancers. FEBS Lett 586, 1602–1605 (2012).

52. Fuhrich, D. G., Lessey, B. A. & Savaris, R. F. Comparison of HSCORE assessment of endometrial beta3 integrin subunit expression with digital HSCORE using computerized image analysis (ImageJ). Anal Quant Cytopathol Histpathol 35, 210–6 (2013).

53. Soyal, S. M. et al. Cre-mediated recombination in cell lineages that express the progesterone receptor. Genesis 41, 58–66 (2005).

54. Farley, F. W., Soriano, P., Steffen, L. S. & Dymecki, S. M. Widespread recombinase expression using FLPeR (flipper) mice. Genesis 28, 106–10 (2000).

55. Masserdotti, G. et al. Transcriptional Mechanisms of Proneural Factors and REST in Regulating Neuronal Reprogramming of Astrocytes. Cell Stem Cell 17, 74–88 (2015).

56. Christianson, J. A. & Gebhart, G. F. Assessment of colon sensitivity by luminal distension in mice. Nat Protoc 2, 2624–2631 (2007).

